# Evaluating and Comparing Measures of Aperiodic Neural Activity

**DOI:** 10.1101/2024.09.15.613114

**Authors:** Thomas Donoghue, Ryan Hammonds, Eric Lybrand, Leonhard Washcke, Richard Gao, Bradley Voytek

## Abstract

Neuro-electrophysiological recordings contain prominent aperiodic activity – meaning irregular activity, with no characteristic frequency – which has variously been referred to as 1/f (or 1/f-like activity), fractal, or ‘scale-free’ activity. Previous work has established that aperiodic features of neural activity is dynamic and variable, relating (between subjects) to healthy aging and to clinical diagnoses, and also (within subjects) tracking conscious states and behavioral performance. There are, however, a wide variety of conceptual frameworks and associated methods for the analyses and interpretation of aperiodic activity – for example, time domain measures such as the autocorrelation, fractal measures, and/or various complexity and entropy measures, as well as measures of the aperiodic exponent in the frequency domain. There is a lack of clear understanding of how these different measures relate to each other and to what extent they reflect the same or different properties of the data, which makes it difficult to synthesize results across approaches and complicates our overall understanding of the properties, biological significance, and demographic, clinical, and behavioral correlates of aperiodic neural activity. To address this problem, in this project we systematically survey the different approaches for measuring aperiodic neural activity, starting with an automated literature analysis to curate a collection of the most common methods. We then evaluate and compare these methods, using statistically representative time series simulations. In doing so, we establish consistent relationships between the measures, showing that much of what they capture reflects shared variance – though with some notable idiosyncrasies. Broadly, frequency domain methods are more specific to aperiodic features of the data, whereas time domain measures are more impacted by oscillatory activity. We extend this analysis by applying the measures to a series of empirical EEG and iEEG datasets, replicating the simulation results. We conclude by summarizing the relationships between the multiple methods, emphasizing opportunities for re-examining previous findings and for future work.

## 1. Introduction

Since the early days of recording electrical activity from the brain, researchers have examined patterns of irregularity, unpredictability, or aperiodicity in neuro-electrophysiological recordings – which we will here refer to as aperiodic activity, as a broad designation for non-rhythmic features of neural recordings. This includes the early observation of an exponential pattern of power across frequencies (Motokawa, 1949), and the use of autocorrelation methods to identify and quantify aperiodic and periodic components (Brazier & Barlow, 1956). Following advances in digital signal processing and the availability of computational analyses, additional methods and conceptualizations came to be applied to neuro-electrophysiological recordings with the goal of characterizing non-rhythmic properties of EEG data, including time domain measures of EEG complexity (Hjorth, 1970), fractal dimension (Nan & Jinghua, 1988), and entropy (Richman & Moorman, 2000), as well as measures of patterns of power in the frequency domain that lack any characteristic frequency (Dumermuth et al., 1977; Kingma et al., 1976; Pascual-Marqui et al., 1988).

Collectively, these early investigations and developments laid the groundwork for ongoing areas of inquiry investigating what can be broadly construed as aperiodic properties of neuro-electrophysiological recordings (Fig 1). However, this research topic is also somewhat fractured, as much of the work has been developed and deployed independently, often engaging distinct and idiosyncratic methods, conceptualizations, and theoretical constructs. This has led to a broad literature in which while there are multiple approaches that share conceptual elements relating to examining ‘aperiodic’ aspects of the data, these different approaches also reflect varying motivations and interpretations. This is reflected in the broad set of distinct concepts and methods that have been employed to study neural data (Fig 2A-B). In addition to these distinct origins and developments, there has thus far been relatively little work to integrate and compare across these approaches. This is despite the increasing attention to studies of aperiodic activity and the increasing prevalence of studies using these methods (Fig 2C), which is split across investigations employing numerous different analysis methods (Fig 2D).

**Figure 1).**
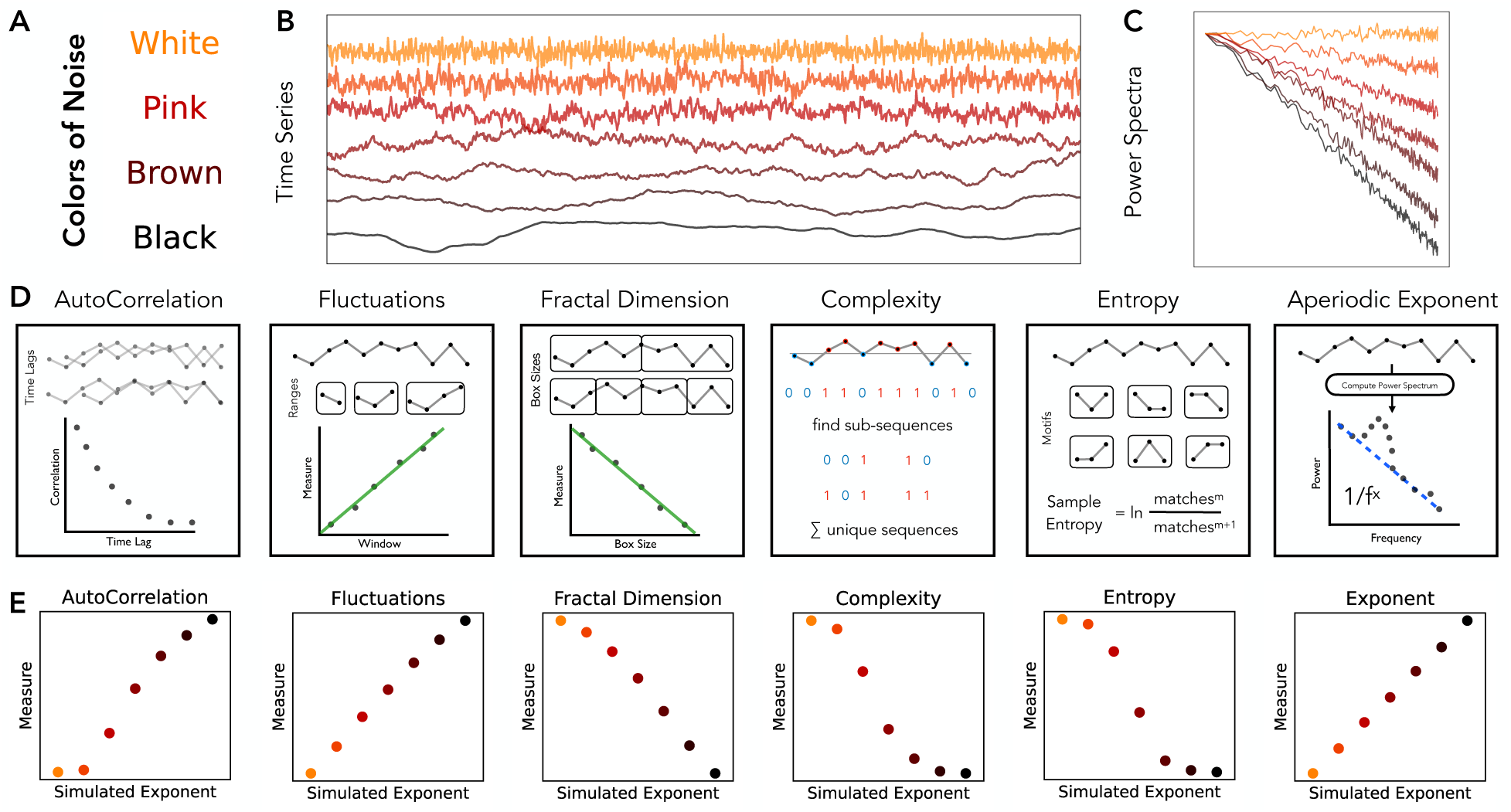
Overview of aperiodic time series and different analysis methods that can be applied to them. **A)** Aperiodic time series can be simulated as “colored noise”, reflecting signals that have a power spectrum with a 1/*f^χ^* distribution, whereby different values of *χ* reflect different named patterns of colored noise: 0 – white; 1 – pink; 2 – brown; >2 – black. Note that for real neural data, X can be equal to non-integer values. **B)** Example simulated aperiodic time series. **C)** Power spectra for the time series in B. **D)** Overview of the kinds of methods that can be applied to time series to characterize aperiodic related features of the data. Methods are grouped together into related categories. D) Result of applying an example method from each category to the example signals from **A.** Empirically, there are clearly similarities in the patterns of results across these signals, however the relationships between these different methods are largely unknown. Note that for each method category, a single example method is applied - specifically autocorrelation decay time (autocorrelation), detrended fluctuation analysis (fluctuations), Higuchi fractal dimension (fractal dimension), Lempel-Ziv complexity (complexity), approximate entropy (entropy), and spectral parameterization (exponent).

Within this broad but disconnected literature there is currently a lack of consensus on methods, interpretations, and best practice guidelines for investigations of aperiodic activity in neural field data. This lack of consensus and integration is salient in relation to the recent emphasis on understanding the different features of neural recordings – whether that be conceptualized as relating to time domain measures of entropy or complexity (Lau et al., 2022) and/or the prominent 1/f-like structure of neural power spectra (He, 2014). In parallel, there have been methodological developments, including work focused on developing methods that can differentiate between aperiodic and periodic components (Donoghue & Watrous, 2023). Given the historical literature and current emphasis on developing an understanding of how different methods relate to each other, and what features of the data they are sensitive to, there is a need for clear and systematic comparisons of extant methods.

This need for a clearer understanding of the relationship between these multiple conceptualizations and measurements of aperiodic neural activity is motivated by examining the findings across the distinct literatures of each approach. For example, aperiodic activity as measured by fractal dimension has been shown to be a dynamic signal that correlates with age (Zappasodi et al., 2015), sleep stages (Ma et al., 2018), anesthesia (Kesić & Spasić, 2016), and task performance (Lutzenberger et al., 1992); whereas aperiodic activity as measured by entropy measures has been shown to be a dynamic signal that correlates with age (Kosciessa, Kloosterman, et al., 2020; Waschke et al., 2017), sleep stages (Miskovic et al., 2019), anesthesia (Liang et al., 2015), and task performance (Sheehan et al., 2018; Waschke et al., 2019); and also aperiodic activity as measured by the spectral exponent has been shown to be a dynamic signal that correlates with age (Donoghue, Dominguez, et al., 2020; Voytek et al., 2015), sleep stages (Ameen et al., 2024; Lendner et al., 2020), anesthesia (Colombo et al., 2019) and task performance (Ouyang et al., 2020; Podvalny et al., 2015; Waschke, Donoghue, et al., 2021), and so on. These patterns of similar findings based on similar measures, examined in distinct investigations that largely do not discuss each other, raise key questions regarding the degree to which these related measures may reflect the same underlying pattern(s) of activity, that are being idiosyncratically measured, reported, and discussed across different subsets of the literature, and/or to what extent these different approaches capture independent variance in the data that can and should be interpreted separately from each other.

**Figure 2).**
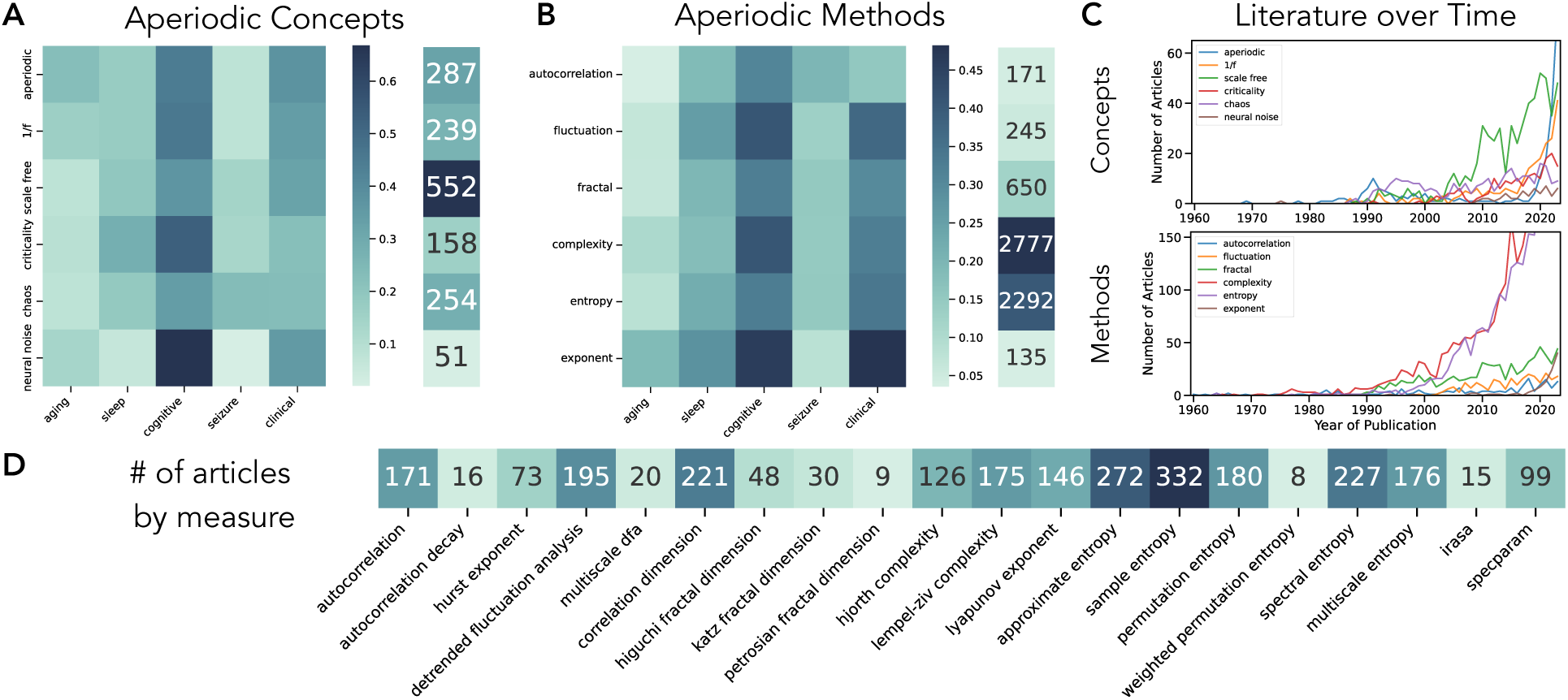
Literature search for aperiodic concepts, methods, and measures in the literature. **A)** Literature search for aperiodic concept terms, and co-occurrence terms from the PubMed database. In the co-occurrence matrix, each cell reflects the proportion of papers with the given aperiodic concept term that co-occurs with the listed term, showing how aperiodic terms are discussed across research topics. The column to the right shows the total numbers of papers identified per concept term. **B)** Literature search for aperiodic method terms, as in A. **C)** Number of aperiodic related articles published across time, separately for ‘concept’ and ‘method’ terms. **D)** Number of articles identified for each individual method included in this project. All literature collections done with the *lisc* Python module. Each of the search terms is identified by a label, whereby each search also used synonyms, inclusion, and exclusion terms to specify literature of interest. Full sets of search terms are available in the project repository.

The use of these different methods to study aperiodicity in neural data draws on developments in time series analysis and digital signal processing (DSP) in engineering and physics. As such, many of the methods and interpretations that have been applied to neural data draw directly from those fields. In doing so, however, it is important to consider that neural field data has some idiosyncratic features that differentiate it from other systems in which analyses of aperiodic signals – such as colored noise or 1/f-related properties – are also common. Neural time series also contain periodic activity (neural oscillations), which are also variable and dynamic within and between subjects (Buzsáki & Draguhn, 2004). Some methods that have been applied to neural data are intended to specifically measure aperiodic properties of the data, being designed to ignore or control for periodic components in the data (Donoghue & Watrous, 2023).

Other methods make no such distinction, and as such may likely reflect a mixture of periodic and aperiodic activity in the context of neural data – even if they were originally designed to measure aperiodic features in other, non-neuroscience related, fields where the data have different properties and generative processes. As such, a clear delineation of which properties of the data methods are sensitive to – and what this means for their interpretation, comparison to other methods, and appropriateness for use in neural data – is also a key consideration when addressing measures of aperiodic neural activity.

Collectively, there is a broad range of literature on aperiodic activity in neural recordings – albeit split and disconnected across different areas of investigation. In current work, it is common to suggest that aperiodic activity is understudied and/or only just emerging as a topic of investigation. However, despite the relative breadth of the literature when considering the different ways of examining aperiodic activity, a lack of clear comparisons and established connections between the various methods and conceptualizations in existing work means that establishing the level of consensus and consistency (or lack thereof) across previous work is quite difficult. To address this issue, this project engages in a systematic evaluation and comparison of a large set of methods ostensibly designed to measure aperiodic activity, identified based on a systematic review of the literature, and compared across simulated data and multiple datasets of extra- and intra-cranial electrophysiological recordings. In doing so, we establish the similarities and differences across this set of methods, providing a putative mapping between the different methods. This mapping can be used to evaluate similarities and differences between the findings and interpretations across the previous literature.

## 2. Methods

To identify methods related to measuring aperiodic activity (broadly construed) in neuro-electrophysiological recordings, we performed a literature analysis. Based on the literature results, we then collected and evaluated multiple methods for estimating aperiodic properties, including time domain methods for autocorrelation, fluctuations, fractal dimension, complexity, and entropy, as well as frequency domain spectral fitting measures. We used simulated data, simulated as either time series or power spectra, that mimic neural data, defined as having aperiodic activity, and in some cases also added periodic components. All simulations and analyses were done using the Python programming language (version 3.9). Time series simulations and analysis methods were done using *neurodsp* (Cole et al., 2019), with some additional methods and utilities also used from *antropy,* for some information theory metrics (Vallat, 2023), *neurokit2*, for some additional complexity measures (Makowski et al., 2021), and *specparam*, for frequency domain simulations and spectral parameterization measures (Donoghue, Haller, et al., 2020). In addition, a series of empirical EEG and iEEG datasets were used to further evaluate the methods under study.

### 2.1 Literature Analysis

First, to evaluate the use of different aperiodic methods, we performed a literature analysis using the Literature Scanner *lisc* Python toolbox (Donoghue, 2019), which allows for collecting and analyzing literature data based on search terms of interest. For this analysis, a list of search terms was curated, including aperiodic methods and concepts, as well as relevant synonyms, and inclusion and exclusion terms which can be used to avoid irrelevant literature. Searches were performed that returned articles with the specified search terms in the titles and/or abstracts of articles indexed in the PubMed database. Initially, systematic searches were used to evaluate the use and prominence of different analysis methods, with methods included if searches returned at least 5 papers including the method in the title of abstract of papers that also discussed neural data analysis, from which we curated a set of method terms, as well as association terms to map these methods to related concepts and topics of inquiry. For co-occurrence analyses, results were normalized by the total number of papers found for a given term (e.g. for co-occurrence of search terms A & B, the normalized score was calculated as count(A & B) / count(A)). For analyses over time, literature searches were run in 1-year increments for the time ranges of 1960 to 2023, inclusive. The full set of search terms, including inclusion and exclusion terms, is available in the project repository.

### 2.2 Overview of Aperiodic Activity & Neural Time Series

The practical definition of aperiodic activity for the simulations and analyses here was based on creating signals with 1/*f^χ^* properties. By 1/f, it is meant that there is a power-law relation between power and frequency, reflecting exponentially decreasing power across increasing frequencies. This consistent pattern of changing power across frequency (regardless of the frequency range) is also referred to as being ‘scale-free’, as the property holds across scales (frequencies). Notably, empirical electrophysiological recordings of neural field data have 1/*f^χ^* - like properties. Different values of *χ* – which we will refer to as the aperiodic exponent – reflect differences in the exponential decay of power across frequencies. Such signals are sometimes also referred to as ‘colored noise’, whereby different colors of noise reflect specific values of *χ*. For example, *χ* of 0 is white noise, which reflects an equal pattern of power across all frequencies, with increasing values of *χ* reflecting steeper patterns of decreasing power across increasing power, whereby *χ* of 1 is known as pink noise, *χ* of 2 as brown noise, and *χ* >2 as black noise. We will refer to this 1/*f^χ^* feature of the data as the aperiodic ‘component’ of the data, with different specifications for *χ* reflecting different variants of the aperiodic component. Note that this 1/*f^χ^* relationship is equivalent to a linear relationship between frequency and power in log-log spacing, such that what is sometimes measured by a line in log-log space, and referred to as the ‘spectral slope’ (b), is equivalent to the aperiodic exponent (*χ* = -b).

Another important aspect of aperiodic activity in neural data is that although neural data has 1/f-like properties, the data are not truly scale-free, 1/f signals: they also have ‘knees’, or frequencies at which the 1/f nature of the signal ‘bends’ (Miller et al, 2009). These ‘knees’ in the data are themselves variable in their occurrence and position (Gao et al., 2020). These ‘knees’ are consistent with multifractal signals, meaning that while they have 1/f properties, there is not a single value of *χ* that describes the power spectrum across all frequencies – rather there are multiple values of *χ*, each of which reflects a particular frequency range, with changes in *χ* occurring at the ‘knee’ points. These ‘knees’ are commonly observed in intracranial data (Gao et al., 2020; Miller et al., 2009), though tend to be less prominently observed in extracranial data (Donoghue, Haller, et al., 2020). The presence of knees in neural data is relevant in considering methods adopted from other areas and whether they pre-suppose a single aperiodic exponent (presuming scale-freeness), and/or allow for what can be called ‘multifractal’ signals (signals with multiple, different 1/f ranges).

In examining neural time series, an additional key consideration is that of neural oscillations, which are distinguished from aperiodic activity by their repetitive, predictable, and rhythmic qualities. Neural oscillations are a common feature and topic of investigation in neural data (Buzsáki & Draguhn, 2004). Notably, oscillatory components have multiple characteristic features, including the center frequency, amplitude, and bandwidth of each oscillation, as well as temporal characteristics such as whether the oscillation is consistent and/or occurs in bursts. Oscillatory activity is by definition rhythmic, occurring at a particular frequency (or scale, thus not being scale-free), and thus has implications for analysis methods, including if and how they attempt to ‘correct’ for oscillatory activity. Here we will refer to signals simulated with both an aperiodic component as well as a periodic component as ‘combined’ signals, which are assumed to be more reflective of actual neural data than purely aperiodic signals. Considering how methods perform on combined signals is important for ecological validity, in particular when considering methods adapted from other areas in which the data under study may not contain periodic components.

### 2.3 Simulations

Time series were simulated to reflect neural data, as combinations of aperiodic and periodic activity. Simulated time series created to reflect power law statistics, with a single 1/f property, were simulated by creating white noise time series, rotating the power spectrum to a specified spectral exponent, and applying an inverse Fourier transform to return to the time domain (Timmer & Konig, 1995). Aperiodic activity displaying a ‘knee’, or a bend in the 1/f, was simulated using a simple physiologically inspired model that combines simulated excitatory and inhibitory post-synaptic potentials, and produces a 1/f-like aperiodic signal with a ‘knee’ (Gao et al., 2017). Simulations including periodic components were simulated by additively combining aperiodic signals with simulated periodic signals, using a periodic kernel, with varying amplitude and frequency characteristics. All time-series simulations were generated using the *neurodsp* toolbox (Cole et al., 2019).

Simulated time series were created to evaluate and compare each of the analyzed methods. All signals were simulated for a length of 30 seconds, at a sampling rate of 250 Hz for the simulations used to test time domain methods, and 500 Hz for testing frequency domain methods. For individual method evaluations, sets of simulations were created to evaluate the effect of each parameter independently, in which each set of simulations systematically varied across a single simulation parameter (Figure 3). Simulations were created to step across aperiodic parameters: aperiodic exponent (range: 0-3 au; increment 0.5), and the aperiodic knee (synaptic decay time sampled from [0.005. 0.015, 0.030, 0.050, 0.075], which systematically alters the knee positions, as simulated with the aforementioned physiological model), and periodic parameters: frequency (range: 5-35 Hz; increment: 1), power (range: 0-2 au; increment 0.1), bandwidth (range: 0.5-3.0 Hz; increment 0.5), and burst probability (range: 0.2-0.8 probability; increment: 0.1). For each parameter value, 50 separate simulations were created. For method comparisons, signals (n=1000) were simulated as either pure aperiodic signals (30%), sampling randomly with exponent values (range: 0-2.5 au; increment: 0.1 au), or as combined signals (70%), sampling the aperiodic component the same way, and adding on a periodic component, sampled with a randomly selected oscillatory center frequency (range: 5-35 Hz; increment: 1) and power (range 0.1-1.0 au; increment: 0.1).

While the majority of the simulations were simulated as time series, for comparisons of a set of spectral fitting methods, power spectra were also directly simulated. Power spectra were simulated to reflect neural power spectra, as combinations of an aperiodic component with overlying peaks. Aperiodic components were simulated with exponential functions describing 1/f forms, including with or without a knee. Periodic components were simulated as Gaussians that were then added on to the aperiodic component. Noise was also added to the power spectra as white noise across all frequencies. All power spectra were simulated using the equations and code described in the *specparam* toolbox (Donoghue, Haller, et al., 2020). Simulations were created with aperiodic exponents of [0.5-3.0, 0.5 increments], with added periodic peaks. Since some methods use an exclusion zone, ignoring frequency ranges that often have oscillations, for these simulations, it was important to mimic the occurrence probability of oscillatory peaks at particular frequencies. To do so, the center frequency of simulated peaks were drawn from the range of 3–34 Hz (1-Hz steps), with each center frequency sampled as the observed probability of center frequencies at that frequency in a large MEG dataset (Donoghue, Haller, et al., 2020). These peaks were sampled with oscillatory power came from the range [0.15, 0.25, 0.5, 1.0, 1.5] a.u. and bandwidth from the range [1.0, 1.5, 2.0, 2.5] Hz, with each value having equal probability.

**Figure 3).**
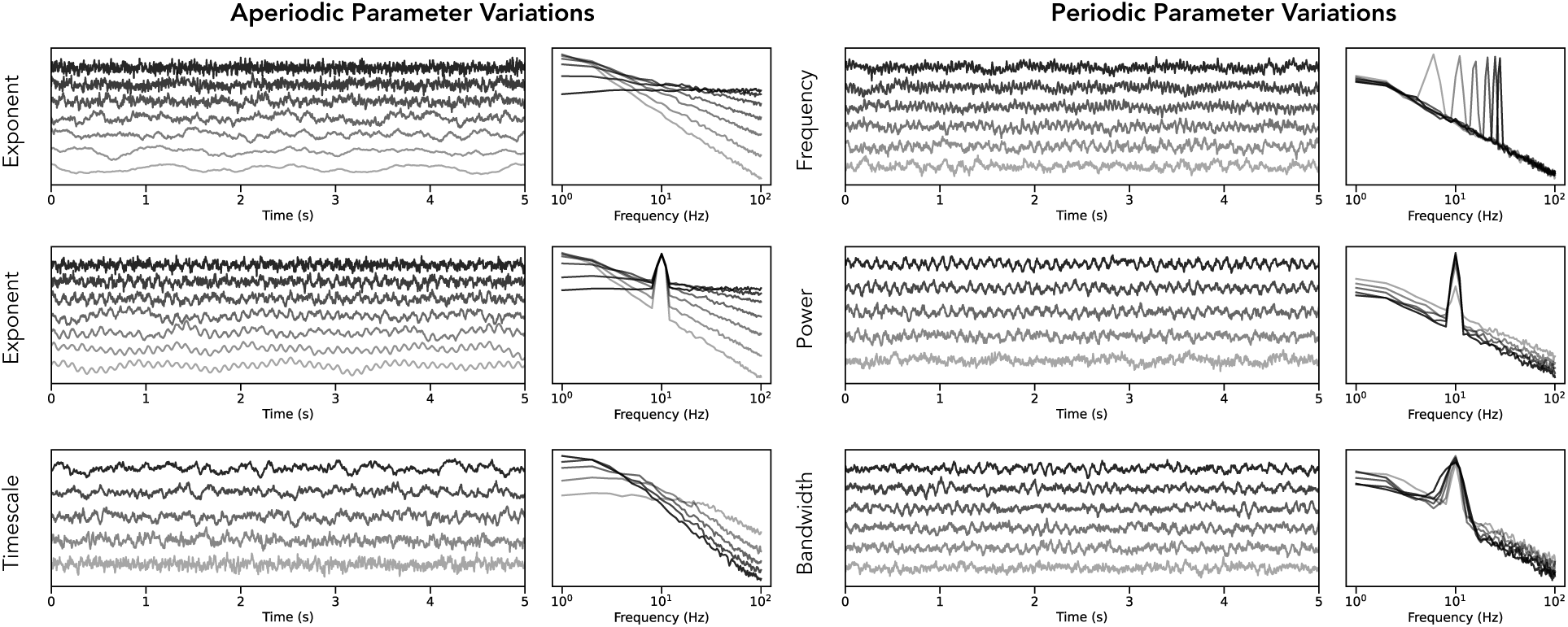
Time series simulations. Time series were simulated across different parameter ranges for both aperiodic (left column) and periodic (right column) parameters. Each panel shows a range of example time series (left) with their corresponding power spectra (right). Examined parameters include varying the aperiodic exponent (left, top), varying the aperiodic exponent in the presence of an oscillation (left, middle), varying the timescale of time series data simulated from a model of post-synaptic potentials (left, bottom), varying the center frequency of an oscillation (right, top), varying the relative power of an oscillation (right, middle), and varying the bandwidth of the oscillatory component (right, bottom). All time series simulations were created with the *neurodsp* Python module.

### 2.4 Time Domain Measures of Aperiodic Activity

Based on the results of the literature analyses, we curated a collection of methods that have been applied to estimate aperiodic features of neural data, starting with methods that are applied in the time domain. We heuristically organized these measures into five groupings – autocorrelation measures, fluctuation measures, fractal dimension measures, complexity measures, and entropy measures. Note that this clustering of methods reflects a practical grouping of related methods, though not a precise technical definition of different types of measures. There are also similarities across the different groups, perhaps most saliently that autocorrelation, fluctuation, and fractal dimension measures are predicated on examining structure across scales (and as such are often interpreted and discussed in similar ways, with reference to the correlation properties or ‘memory’ of signals). The measures grouped as complexity and entropy measures are more typically conceptualized in relation to reflecting the variability of the signal. In the following we describe a broad set of methods identified through this project. The analyses presented in this paper are restricted to a set of highlighted methods, with evaluations of the remaining methods also available in the project repository (Table 1). Notably, we focus here on popular mono-scale time-domain methods, without focusing on multi-scale methods or methods that first require constructing a state space. Settings for each method are reported in Table 1.

**Table 1:**
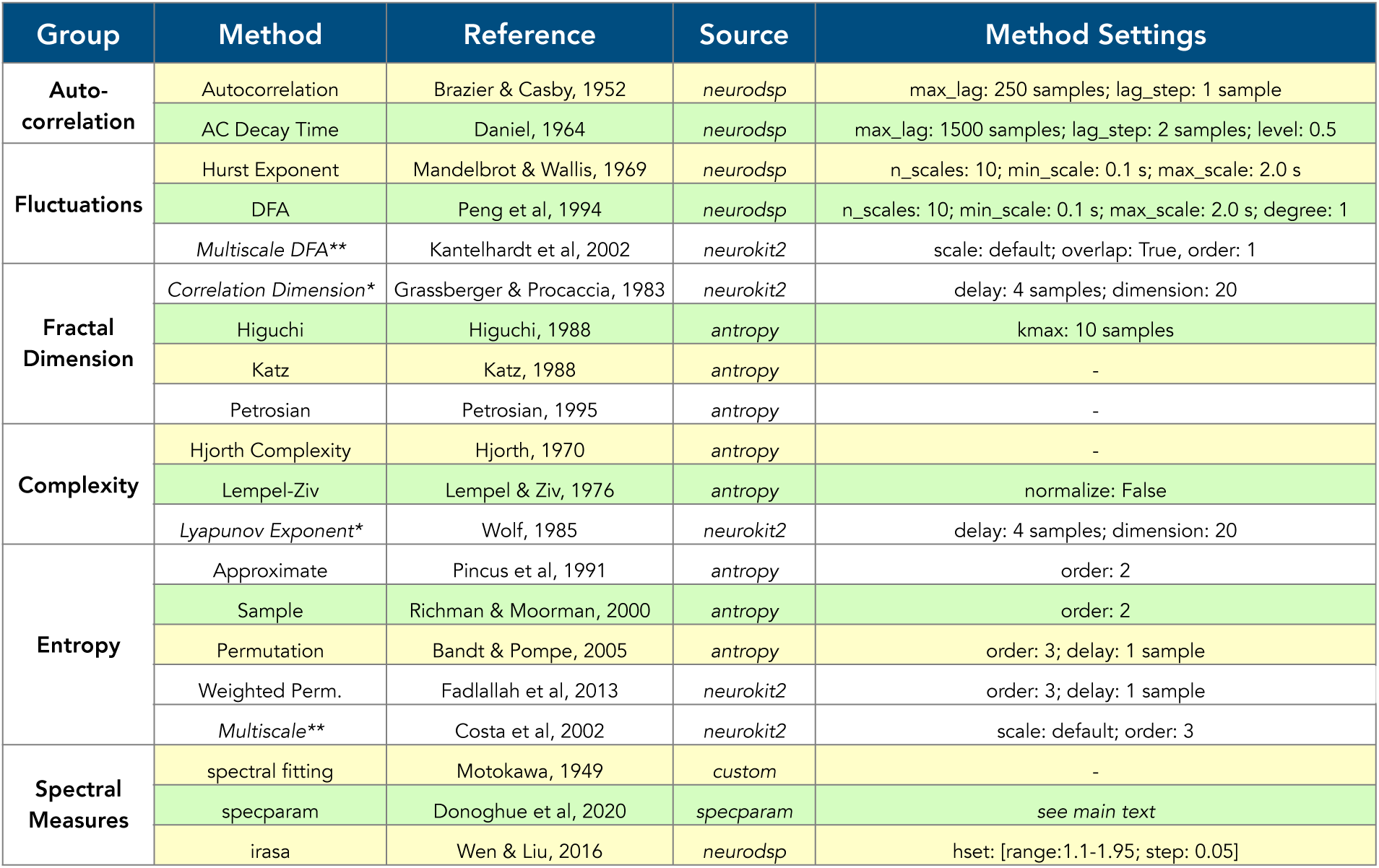
Aperiodic Methods. Each method is listed with its group affiliation, original reference, and the source of the implementation and methods settings used. Methods in green are used as example methods for the category, chosen based on occurrence in the literature, and are featured throughout the manuscript. Methods in yellow are also included in analyses in some figures. Methods in white are additional methods not reported in the manuscript either, including those that were excluded as they require the computation of a state space (marked with *) or due to being multi scale methods (marked with **). Full results for all methods (including in white) are available in the project repository. Abbreviations: AC: autocorrelation; FD: fractal dimension; DFA: detrended fluctuation analysis; s: seconds.

| Group | Method | Reference | Source | Method Settings |
| --- | --- | --- | --- | --- |
| Auto-correlation | Autocorrelation | Brazier & Casby, 1952 | neurodsp | max_lag: 250 samples; lag_step: 1 sample |
|  | AC Decay Time | Daniel, 1964 | neurodsp | max_lag: 1500 samples; lag_step: 2 samples; level: 0.5 |
| Fluctuations | Hurst Exponent | Mandelbrot & Wallis, 1969 | neurodsp | n_scales: 10; min_scale: 0.1 s; max_scale: 2.0 s |
|  | DFA | Peng et al, 1994 | neurodsp | n_scales: 10; min_scale: 0.1 s; max_scale: 2.0 s; degree: 1 |
|  | Multiscale DFA** | Kantelhardt et al, 2002 | neurokit2 | scale: default; overlap: True, order: 1 |
| Fractal Dimension | Correlation Dimension* | Grassberger & Procaccia, 1983 | neurokit2 | delay: 4 samples; dimension: 20 |
|  | Higuchi | Higuchi, 1988 | antropy | kmax: 10 samples |
|  | Katz | Katz, 1988 | antropy | - |
|  | Petrosian | Petrosian, 1995 | antropy | - |
| Complexity | Hjorth Complexity | Hjorth, 1970 | antropy | - |
|  | Lempel-Ziv | Lempel & Ziv, 1976 | antropy | normalize: False |
|  | Lyapunov Exponent* | Wolf, 1985 | neurokit2 | delay: 4 samples; dimension: 20 |
| Entropy | Approximate | Pincus et al, 1991 | antropy | order: 2 |
|  | Sample | Richman & Moorman, 2000 | antropy | order: 2 |
|  | Permutation | Bandt & Pompe, 2005 | antropy | order: 3; delay: 1 sample |
|  | Weighted Perm. | Fadlallah et al, 2013 | neurokit2 | order: 3; delay: 1 sample |
|  | Multiscale** | Costa et al, 2002 | neurokit2 | scale: default; order: 3 |
| Spectral Measures | spectral fitting | Motokawa, 1949 | custom | - |
|  | specparam | Donoghue et al, 2020 | specparam | see main text |
|  | irasa | Wen & Liu, 2016 | neurodsp | hset: [range:1.1-1.95; step: 0.05] |

One of the categories of methods that have long been applied to EEG data to measure (a)periodicity are autocorrelation measures. The signal autocorrelation reflects the correlation of the signal to itself, with a certain time-lag, with this autocorrelation typically measured across different time lags. Beginning from the early days of computational analyses, measures of autocorrelation were applied to quantify aperiodic and periodic components (Brazier & Barlow, 1956; Brazier & Casby, 1952). Similarly, ratio measures were designed to capture the ratio of the dominant rhythm over the background activity (Daniel, 1964) and early analyses showed that task activation engaged an increase in aperiodic components of the EEG activity (Matoušek et al., 1969). In this investigation, we computed the autocorrelation function across a set of time lags as well as the ‘autocorrelation decay’, which is a measure of the minimal time lag for the autocorrelation function to decay to a specified value (here, autocorrelation of 0.5), which has also been applied to neural recordings (Gao et al., 2020; Honey et al., 2012).

Related to the autocorrelation, a set of methods we will here refer to as ‘fluctuation’ methods investigate how properties of the data vary across different window sizes. For example, rescaled range measures the variability of time-series across different sized segments of the data, from which the Hurst exponent can be calculated (Mandelbrot & Wallis, 1969). The Hurst exponent reflects a measure of what is sometimes referred to as the ‘long-term memory’ of a time series – reflecting a measure of the rate of decay of statistical dependence across different time ranges. Similarly, detrended fluctuation analysis (*dfa*), first developed for applications in genetics (Peng et al., 1994), computes local linear fits across window sizes as a measure of the pattern(s) of temporal autocorrelation in the signal. This method was subsequently applied to EEG data (Ferree & Hwa, 2003; Watters & Martin, 2004).

For fluctuation measures, while these measures are relatively common in neuroscience, how they are applied does vary such that not all applications are conceptually consistent with the topic here of analyzing aperiodic variation in recorded neural field data. Notably, *dfa* in particular is often applied to analyze the properties of the amplitude envelope of narrowband filtered time series (Hardstone et al., 2012; Linkenkaer-Hansen et al., 2001). This approach is used to evaluate and interpret long-term correlations in the amplitude variations of narrowband, putatively oscillatory, frequency ranges, which we consider a distinct application and interpretation as to what is being investigated here. In this work, we computed the Hurst exponent and the alpha value from *dfa*, as measures of the broadband signal, meaning they were computed from the original (non-narrowband filtered) time series, and all references to *dfa* and related literature refer to this kind of application, unless explicitly noted.

Another set of methods related to fluctuations measures are fractal dimension measures, which are measures of the complexity of self-similar patterns (Mandelbrot, 1967), whereby fractal refers to entities with the same or similar structure across different scales. Multiple variants of fractal dimension measures have been applied to neural time series (Accardo et al., 1997; Esteller et al., 2001; Kesić & Spasić, 2016; Nan & Jinghua, 1988). These fractal dimension measures are based on the ‘box counting’ dimension, which estimates the number of entries in a grid needed to cover a shape (or signal), across different grid sizes. They estimate the curve length of the signal across different signal lengths, with the corresponding dimension being the best-fitting slope of the log-log plot of curve lengths over signal lengths. In this investigation we employed several measures of the fractal dimension that have been applied to EEG data, including Higuchi (Higuchi, 1988), Katz (Katz, 1988), and Petrosian (Petrosian, 1995).

Distinct from methods based on correlation and signal properties across signal lengths are measures that are broadly construed to measure the variability of neural time series. We first consider a collection of methods that we will refer to as ‘complexity’ measures – though note that while this reflects a conceptual similarity in the goals of such methods, they do not entail a specific or singular mathematical or methodological approach for defining and measuring complexity. We include in this category the Hjorth parameters, a set of parameters designed to describe EEG data, including Hjorth activity, Hjorth mobility, and Hjorth complexity (Hjorth, 1970). In addition, Lempel-Ziv complexity characterizes the complexity of a time series based on the number of sub-strings encountered in binary sequence (Lempel & Ziv, 1976), which can be applied to EEG data by binarizing recorded voltage values (Medel et al., 2023; Zhang et al., 2001).

Finally, similar to complexity measures, we include a set of entropy measures, based on information theory, that have also been developed and applied to time series data as measures of signal variability / complexity. The use of these various entropy measures is quite common in neuroscientific investigations (Lau et al., 2022). The included time domain measures of entropy are approximate entropy, which quantifies the amount of regularity in time series based on estimating the likelihood that patterns of data remain similar on incremental comparisons (Pincus et al., 1991), the updated variant, sample entropy, which similarly quantifies regularity while being more robust to signal length (Richman & Moorman, 2000), permutation entropy, which characterizes the signal based on the frequency of occurrence of different motifs in the signal (Bandt & Pompe, 2002), and a variation, weighted permutation entropy, which is updated to weight motifs by the variance of the data segments (Fadlallah et al., 2013). Entropy measures can also be also be applied to frequency domain representations (spectral entropy), which is also included in the project repository (Inouye et al., 1991).

In collecting and evaluating methods for this project, we also encountered a series of methods that draw from dynamical systems and/or chaos theory, which include constructing a phase (or state) space of the signal and then computing measures upon this representation. A phase space is a representation of the states of a dynamical system in which all the states of the system are represented. In this project, we included some such methods in the literature search and project repository, but excluded them from the main analyses, due to considering the requisite treatment of dynamical systems and chaos theory in order to be able to productively discuss the assumptions, interpretations, and implications of such measures (as well as their considerably greater computational complexity) to be beyond the scope of the current project. Relevant methods include the correlation dimension, generally considered to be a measure of fractal dimension which measures the dimensionality of the space of a set of points (Grassberger & Procaccia, 1983) that has been examined and interpreted in EEG data (Lutzenberger et al., 1992; Nan & Jinghua, 1988), as well as Lyapunov exponents, generally considered as a measure of complexity, which is a measure from non-linear dynamics that measure the rate of divergence of trajectories in phase-space (Wolf et al., 1985) that has also been applied to EEG data (Babloyantz & Salazar, 1985; Mayer-Kress & Holzfuss, 1987). For reviews of nonlinear measures and their application to EEG data (including discussions of their applicability) see the following reviews and evaluations (Krakovská & Štolc, 2008; Le Van Quyen et al., 2003; Pritchard, 1992; Stam, 2005).

### 2.5 Frequency Domain Measures of Aperiodic Activity

Since 1/f properties can be visualized in frequency domain representations, several related approaches have been developed and applied for measuring 1/f properties directly from power spectra. This includes applying a linear fit using a simple linear regression (Dumermuth et al., 1977; Freeman & Zhai, 2009; Inouye et al., 1994), as well as approaches that have applied exponential, polynomial, or t-distributions to model more variable patterns of power (Dehghani et al., 2010; Kingma et al., 1976; Pascual-Marqui et al., 1988). A key difficulty of measuring aperiodic activity from the power spectrum directly is how to be robust to periodic components such as neural oscillations that exhibit as ‘bumps’ in the power spectrum over and above the aperiodic component. To avoid oscillatory regions, some approaches have fit a linear fit excluding frequency ranges that typically contain oscillatory power (Kosciessa, Grandy, et al., 2020; Voytek et al., 2015). A more generalized approach parameterizes the neural power spectrum, fitting both the aperiodic component and any overlying periodic peaks (Donoghue, Haller, et al., 2020).

To test these variations for measuring the aperiodic component, we employed a series of spectral fitting methods designed to estimate the aperiodic exponent from the power spectrum. This included testing several proposed variants of this approach including approaches to fit a linear fit of the power-spectrum, in log-log, done, as ordinary least squares (OLS) fit, a robust linear model (RLM) fit, and using the RANSAC robust regression algorithm. We also tested an exponential fit (EXP) of the power spectrum in semi-log space, fit as non-linear least squares curve fitting procedure (scipy.optimize.curve_fit). All of the above methods were also fit using oscillation exclusions, as has been done in previous work, excluding a fixed alpha region from fitting to avoid the prominent oscillatory peak (Voytek et al., 2015). We also applied the *specparam* (formerly *fooof*) tool for parameterizing neural power spectra (Donoghue, Haller, et al., 2020), which is itself an adapted method for spectral line fitting. Briefly, this approach seeks to jointly model the 1/f with an exponential as well as modelling overlying oscillatory peaks, fit as Gaussians. It uses an iterative procedure to fit and remove peaks, allowing for a final fit of the 1/f which is fit on a peak-removed version of the original spectrum.

An alternate strategy for measuring aperiodic activity from the power spectrum while controlling for periodic activity is to use coarse-graining (resampling) approaches to isolate the aperiodic component of the spectrum. Since periodic activity has characteristic frequencies, up- or down-sampling the time sampling can displace periodic activity, which can be leveraged to isolate scale-free properties, which are not manipulated by the resampling. An original algorithm for this approach, called coarse-graining spectral analysis (Yamamoto & Hughson, 1991, 1993) was originally proposed for electrocardiography data, but was subsequently applied to neurophysiological data (He et al., 2010). An adapted approach, called Irregular Resampling Auto-Spectral Analysis (*irasa*), was more recently developed and proposed for neural data (Wen & Liu, 2016). In this investigation, we applied the *irasa* method.

For all spectral methods, there are hyperparameters including the selecting the frequency range to fit, as well as selecting the model form to fit to the spectrum. Note that *irasa* is a decomposition method, and not inherently a model fitting method, although models can be fit to the isolated aperiodic component isolated by use of *irasa*. For all comparisons between spectral parameterization and *irasa*, equivalent models were fit to the isolated aperiodic components. For simulations in which data was simulated with a single 1/f component (e.g. colored noise), aperiodic activity was measured on the frequency range of 1-50 Hz, with a 1/*f^χ^* model – which is the ‘fixed’ model in *specparam*, and equivalent to fitting a linear fit in log-log space, with additional *specparam* settings: (max_n_peaks: 8; peak_width_limits: [1, 8]; peak_threshold: 2; min_peak_height: 0.05). In simulations in which data was simulated to include an aperiodic knee, models were fit to the frequency range of 1-100 Hz with a model of the form 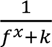 which is the ‘knee’ model from *specparam*, with additional settings: (max_n_peaks: 12; peak_width_limits: [1, 8]; peak_threshold: 2; min_peak_height: 0.1).

### 2.6 Known Relationships between Methods

The above set of methods draw from multiple overlapping yet often distinct traditions of analyses, with variable available information on the expected analytical relationships between methods. In some cases, the theoretical relationship between different measures is known. For example, for colored noise signals, the *dfa* alpha (*α*) value relates to the aperiodic exponent as *α* = (-*χ* + 1) / 2 (Kiyono, 2015; Robinson, 2003; Schaefer et al., 2014), the fractal dimension (D) relates to the *dfa* alpha value as FD = 3 – *α* (Eke et al., 2002; Esteller et al., 2001), and the fractal dimension maps to the aperiodic exponent as D = (5 – *χ*) / 2 (Cervantes-De la Torre et al., 2013; Higuchi, 1990). For detailed discussions on the relationships between these measures, see the following general reviews (Eke et al., 2002; Schaefer et al., 2014). As far as we are aware, there are no analytical solutions to the expected value of the examined complexity or entropy measures on colored noise signals, nor any theoretical expectations for the results of any of the employed time domain methods on combined signals. Where there is a known theoretical expectation of a measure, this is included in the plotted results, based on the relationships noted above.

### 2.7 Empirical Datasets

To evaluate whether the examined relationships between methods hold in empirical datasets, we additionally analyzed EEG and iEEG datasets. An initial dataset comprised eyes closed resting state EEG data collected in the VoytekLab at UC San Diego, recorded on a 64 channel BrainVision system, downsampled to 500 Hz sampling rate for analysis. From this dataset, we extracted 30 second segments of resting state data per subject (n=29, ages=18–28 [mean: 20.62; standard deviation: 1.80], number of males=11). Additionally, we analyzed EEG data from the openly available Multimodal Resource for Studying Information Processing in the Developing Brain (MIPDB) released by the Child Mind Institute (Langer et al., 2017). This dataset includes children and adults (n=126, ages=6–44 [mean: 15.79; standard deviation: 8.03], number of males=69). For this analysis, we used only resting state EEG data which was collected on a 128 channel Geodesic Hydrocel system sampled at 500 Hz, from which we extracted and analyzed the first eyes-closed resting state segment, lasting 30 seconds. The outermost channels, around the chin and neck, were excluded, leaving a standard 111 channel setup. Fifteen subjects were lacking resting state recordings or did not have sufficient data to analyze, such that there were 111 participants included in the analysis of this dataset.

We also analyzed iEEG data from the MNI open iEEG atlas, which contains one-minute recordings from 1772 channels of recordings of quiet, eyes-closed wakefulness (n=106, ages=13–62 [mean: 33.46; standard deviation: 10.67], number of males=58). This dataset is available with preprocessing already applied, including a bandpass filter from [0.5-80 Hz], and having been downsampled to a sampling rate of 200 Hz (Frauscher et al., 2018). This dataset is organized into a set of 38 brain regions, each containing recordings from 5 patients. For this analysis we analyzed only cortical contacts, keeping 1479 channels of data. Details of the dataset preprocessing are available in the dataset description (Frauscher et al., 2018). We extracted 30 seconds from the available dataset, across all channels, and used them for analysis.

For all EEG and iEEG analyses, each method was applied separately to each channel across each subject. The same method settings as used and reported for the simulation tests were used on the empirical data. In addition to the aperiodic measures, we also estimated periodic power, computed as the dominant (highest power) peak as detected by the spectral parameterization method, computed for the alpha range (7-14 Hz) for the EEG datasets, and across the range of 2-35 Hz in the iEEG datasets. For all the EEG and iEEG data, correlations across electrodes were computed by comparing across electrodes – for the first EEG dataset, this was done at Oz, for the second EEG dataset this was done at Cz, and for the iEEG dataset, this was done across all channels together. For the EEG datasets, topographies were computed by averaging each measure across channels across all subjects. Spatial correlations across electrodes were then computed by correlating the average measures across channels. For the EEG datasets, frequency domain measures were computed across the range of 3-40 Hz, and spectral fits used a single exponential (equivalent to a linear fit in log-log or ‘fixed’ mode in *specparam*). In the iEEG data, in which there was evidence of a knee, we computed the spectral fits across the frequency range of 1-60 Hz fit with a knee model. For the *specparam* models, the mean R^2^ was 0.972 for the first EEG dataset, 0.977 for the second EEG dataset, and 0.984 for iEEG dataset, all of which reflect good models fits.

### 2.8 Statistical Comparisons

Each included measure was evaluated across the same set of simulation parameters, to evaluate how each varies across different parameters. For individual method evaluations, measures were computed across parameter variations. Where an expected value was known (e.g. measuring the aperiodic exponent for simulations with a specified aperiodic exponent), errors were computed, with two sample t-tests and Cohen’s d used to compare significant differences and effect sizes between distributions of errors. For pairwise comparisons between measures, correlations were computed between measure results, using Spearman correlations. For all correlation measures, we used bootstrapping approaches to compute confidence intervals (CIs) for correlation coefficients, as well as to test for significant differences between correlation magnitudes (Wilcox, 2016). Bootstrap procedures were calculated with 5000 resamples, computing a distribution of values from which 95% CIs were computed. Bootstrapping procedures were also used to compute and test for differences between correlations, computing a two-sided empirical p-value for a test for a significant difference from 0 from the resampled distribution. To compare the relationships between aperiodic measures and features of interest, such as age in the second EEG dataset, we computed both the Spearman correlations of each measure to age (at electrode Cz), as well as the semi-partial correlation of age to each measure, after regressing out the estimated aperiodic exponent. Differences between full and semi-partial correlations were tested with the bootstrapping procedure described above.

## 3. Results

In this project we systematically evaluated multiple methods that have been applied to investigate aperiodic properties in neuro-electrophysiological recordings (Figure 1). The set of included methods was informed by systematic literature searches that helped to evaluate the use of the different methods that have been employed to analyze aperiodic activity in neural data (Figure 2). The literature analysis demonstrated that there are multiple different conceptualizations of aperiodic neural activity (Figure 2A) across different kinds of methods (Figure 2B). Notably, papers discussing concepts and methods relating to aperiodic activity are increasing over time (Figure 2C). Based on literature search that identified numerous individual methods that have been applied to neural data (Figure 2D), we curated a collection of methods, grouped into autocorrelation, fluctuation, fractal dimension, complexity, entropy, and spectral fitting methods (Table 1).

To evaluate each method on signals for which ground truth parameters are known, we first used a simulation procedure to create time series signals that varied across different parameters of interest (Figure 3). Specifically, simulations were created that systematically varied across aperiodic parameters, including the aperiodic exponent, with and without an overlying oscillation, and across the timescale from a simulation model based on post-synaptic potentials (Gao et al., 2017). In addition, to test the impact of combined signals (including an oscillation), we examined simulations varying across oscillation center frequency, relative power, and bandwidth. Note that due to the large number of methods and simulation parameters included overall, for the sake of space we focus on a specified set of main methods (Table 1) and simulation parameters (variations in the aperiodic exponent, with and without oscillatory components, as well as the effects of oscillatory frequency and power) in the results that are directly reported in this manuscript. Additional methods and simulation tests available in the project repository (https://github.com/AperiodicMethods/AperiodicMethods).

In the first set of method evaluations, we sought to examine the performance of various time-domain methods that have previously been applied to neural data, including autocorrelation, fluctuation, fractal dimension, complexity, and entropy measures on the simulated data (Figure 4). Overall, all the methods display the expected variation across simulated aperiodic activity – following the general interpretation that going from white noise to black noise reflects increasing regularity / decreasing randomness (increasing decay rate, decreasing fractal dimension / complexity / entropy). Notably, however, the relationships between each method’s outputs and the examined simulated parameters typically do not follow a simple, linear relationship – different methods have different relationships with the simulation parameters. Overall, this supports that despite the broad similarity across the different methods, there is also a level of idiosyncrasy across the different methods.

**Figure 4).**
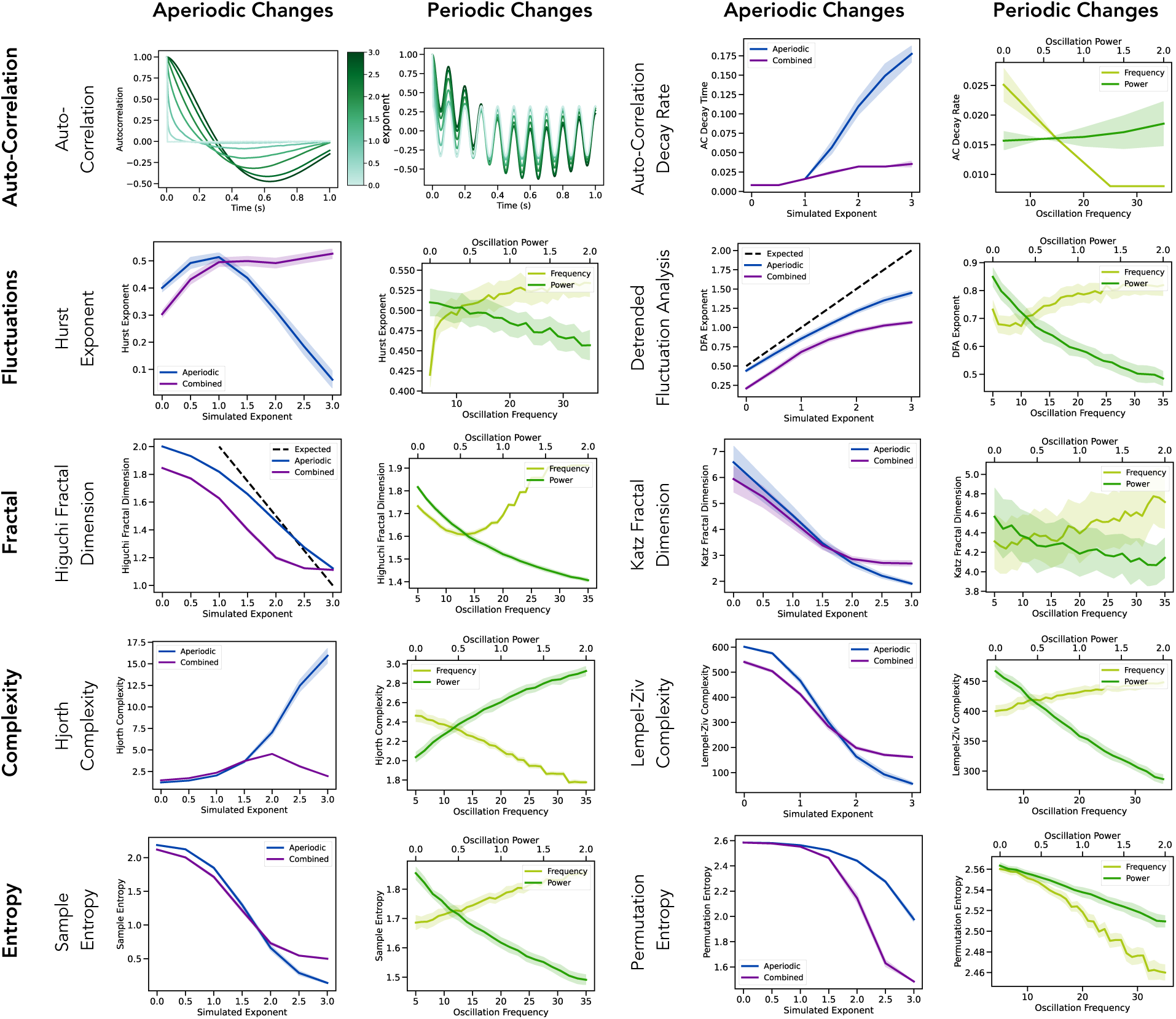
Evaluations of time domain aperiodic measures. Each row reflects a method category and includes two example methods from this category. Columns evaluate the methods on simulated data with variations in either aperiodic or periodic features, whereby ‘aperiodic’ labelled signals reflect variation in the aperiodic component of the signal, ‘combined’ reflect variation of the aperiodic component in signals that also have periodic components, and ‘frequency’ and ‘power’ reflect variation in the center frequency and power of the periodic component. Where available, the expected result of the given measure is indicated in the dashed black line. Notably, all included methods show marked variation across aperiodic parameters, though the specifics of how they do so, and the impacts of periodic components, vary.

Examining the effects of oscillatory features is also consistent with the general notion that periodic activity reflects a degree of regularity to the data, such that increasing oscillatory power generally leads to results that can be interpreted as ‘less irregular’. Again, there are clear differences in these patterns across the different methods. For example, comparing between aperiodic and combined signals highlights some notable patterns, reflecting differences in the extent to which the methods are sensitive to oscillatory components in the data. Some methods (e.g. Lempel-Ziv complexity, Katz fractal dimension, and sample entropy) show only small differences, suggesting these methods are specifically sensitive to the aperiodic characteristics of the data. In contrast, other methods (e.g. Hurst exponent, Hjorth complexity and permutation entropy) show a notable difference between the two cases, reflecting that these methods are more sensitive to oscillatory activity. This is broadly consistent in the results of the simulations across different oscillatory features. By splitting out different oscillatory features, these simulations also show that some methods (Hurst exponent, Katz fractal dimension, permutation entropy) have similar responses to increases in oscillatory frequency and power, whereas other methods (detrended fluctuation analysis, Lempel-Ziv complexity, sample entropy) show opposite parameters, changes in oscillatory activity tend to have smaller impacts on the method results than changes in the aperiodic parameters, supporting the idea that these methods are likely to typically reflect the aperiodic component of the data. However, given that most of the methods are significantly impacted by oscillatory components, they cannot be said to be aperiodic specific measures.

In a subsequent set of analyses, we compared different approaches that have previously been employed to measure aperiodic components from neural power spectra. This set of methods includes basic fitting methods, spectral parameterization, and the *irasa* method, evaluated across simulated aperiodic signals, combined signals, and aperiodic signals with a knee (Figure 5A). In evaluating the spectral fit methods, we found that the spectral parameterization method significantly outperformed the other fitting methods (OLS, Robust linear fit, RANSAC, and exponential fit), including when using an alpha exclusion region (Figure 5B). The spectral parameterization method was robust in estimating the aperiodic exponent on aperiodic signals, combined signals, knee signals, and across variable peak bandwidths (Figure 5C). By comparison, while the *irasa* method excelled at aperiodic and combined signals, estimates were significantly impacted by the presence of a knee, or variable bandwidth peaks (Figure 5D). This is consistent the *irasa* method’s assumptions of purely fractal data, such that it has trouble with data that violates this specific assumption, such as in cases in which there is a knee (Donoghue, Haller, et al., 2020; Gerster et al., 2022; Wen & Liu, 2016). In direct comparisons between *specparam* and *irasa*, the results between the measures tended to be highly correlated, with *irasa* slightly outperforming spectral parameterization on aperiodic-only or combined signals with a sinusoid, where as *specparam* outperformed *irasa* on knee signals or signals with an oscillatory component with a broader bandwidth (Figure 5E).

**Figure 5).**
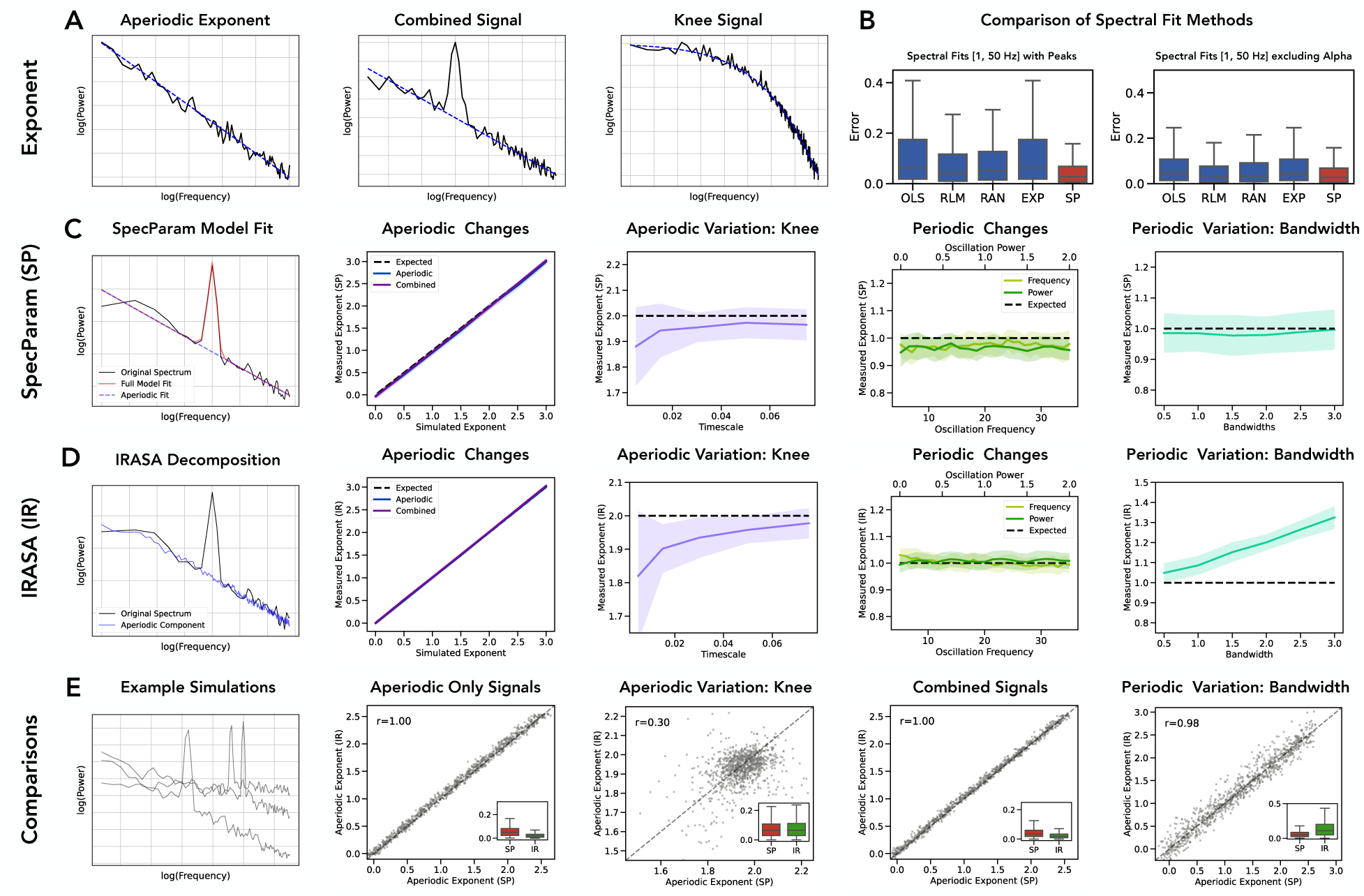
Evaluation and comparison of spectral domain aperiodic measures. **A)** Example simulated power spectra, with the corresponding aperiodic component shown in blue, including a purely aperiodic signal (left), a power spectrum including a periodic component (middle), and a power spectrum with a ‘knee’. **B)** Comparison of measures of the aperiodic exponent done by directly fitting the power spectrum, applied to simulated combined signals (aperiodic components with peaks). **C)** Evaluation of the *specparam* method for measuring aperiodic components, including an example of the method (first panel; left side), an evaluation of performance on aperiodic and combined signals (second panel), an evaluation of the method on ‘knee’ signals (third panel; middle), an evaluation of the method with periodic changes of varying frequency and power (fourth panel) and an evaluation on signals of varying periodic bandwidth (fifth panel; right side). **D)** Evaluation of the *irasa* method, with the same organization as C. E) Direct comparisons of specparam and *irasa* on simulated data. The left panel shows the power spectra of example simulations (combined signals), with subsequent panels following the organization of **C & D.** Insets show the distribution of errors for each method. Abbreviations: OLS: ordinary least squares; RLM: robust linear model; RAN (*ransac*): random sample consensus; EXP: exponential fit. SP (*specparam*): spectral parameterization; IR (*irasa*): irregular resampling auto-spectral analysis.

Having explored each individual method, we then directly compared the methods to each other (Figure 6). We did so by creating simulated data of aperiodic signals and combined signals and applying each of the methods to these simulations. We then examined the methods in a pairwise fashion, comparing the method results and computing the correlations across each combination of methods. For a prioritized subset of the time domain methods, as well as spectral parameterization as a representative of spectral domain methods, we visualized the relationship between the method estimates (Figure 6A), as well as summarizing the correlations between methods on aperiodic-only signals (Figure 6B) and on combined signals (Figure 6C). Overall, this analysis emphasizes that these measures largely covary, in particular for aperiodic only signals (in which the correlation magnitudes are all between 0.97-1.00). However, consistent with the earlier simulations, the methods are differently sensitive to signals with an oscillatory component, with much more variability in such signals. In combined signals, the direction of all the correlations is the same as aperiodic only signals, but the magnitudes range from 0.31-0.95. Notably, the methods are only mildly correlated with the simulated peak power of the oscillation (r-values of 0.02-0.18), suggesting the methods are overall largely sensitive to aperiodic features of the data.

**Figure 6).**
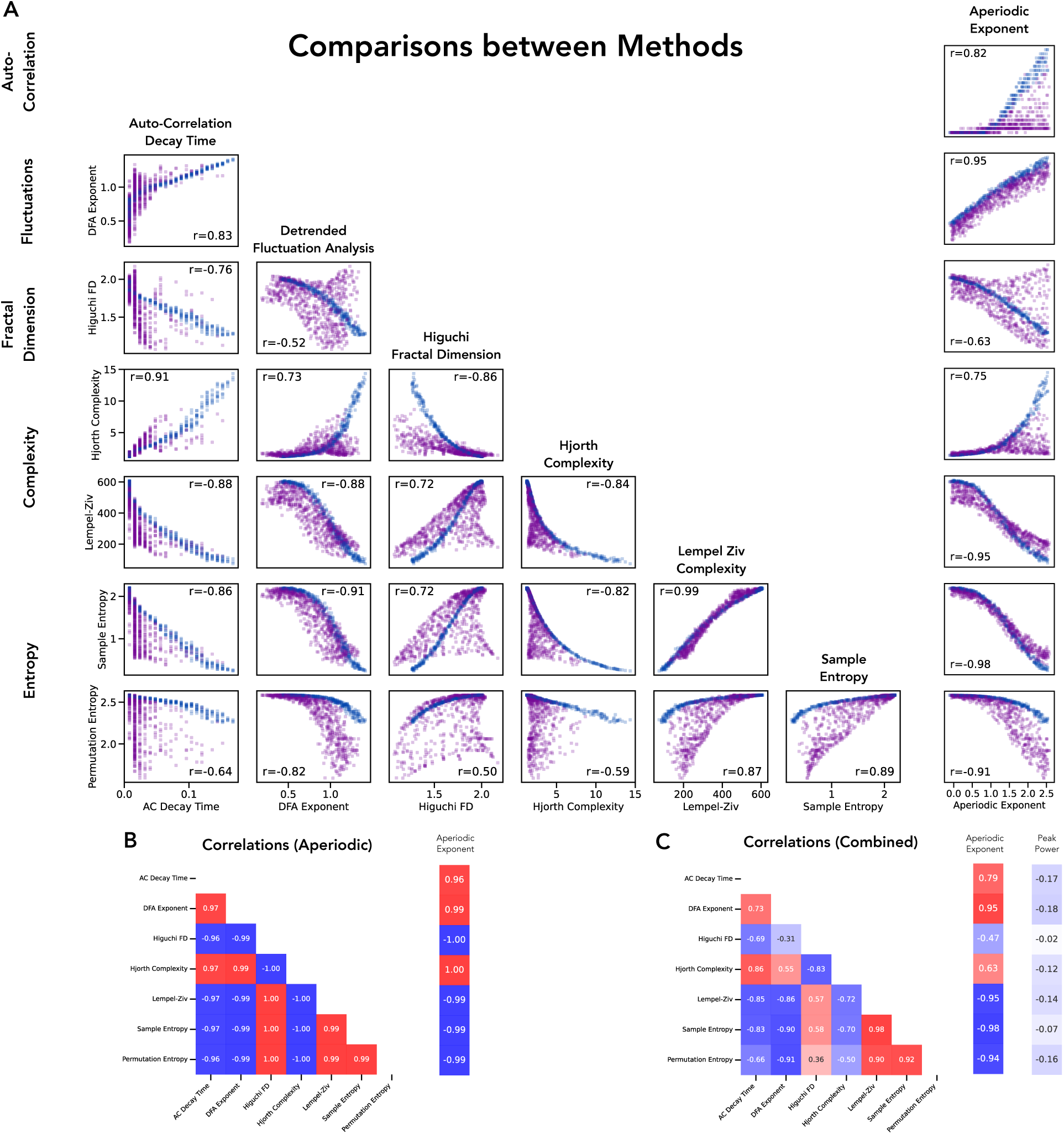
Comparisons between methods, including comparing time domain measures to each other, and to spectral exponent estimation. **A)** Each panel shows the relation between estimates of two methods, with each point demonstrating the measure results for the two methods on a single simulated time series. Time series were simulated across varying aperiodic parameters (blue dots) as well as with varying aperiodic and periodic parameters (purple dots). Insets report the correlation between the measures across all simulations. Notably, there are systematic relationships between all pairs of methods, though with some variation that largely reflects the presence or absence of periodic components. **B)** Correlation matrix comparing measures to each other, for the aperiodic only signals. **C)** Correlation matrix comparing measures to each other, for the combined signals, including comparison to peak power.

We next sought to examine the performance of the aperiodic methods in empirical datasets, to replicate the results from the simulations on empirical data. We started with an initial dataset of eyes closed resting-state EEG data collected from healthy young adults (Fig 6A-B). We first compared the measures of the aperiodic exponent between *specparam* and *irasa* (Fig 6C-D) finding a high correlation, computed both across subjects (r=0.946 [0.854-0.978], p<0.001) and across electrodes (r=0.972 [0.937-0.985], p<0.001). We also computed the dominant oscillatory activity (Fig 6E), as well as a set of time domain methods. We computed the correlations between each time domain measure and the aperiodic exponent (Fig 6F), as well as the spatial topography of each measure (Fig 6G). We computed the correlations between the aperiodic measures, as well as to the periodic measure, across subjects (Fig 6H) and across electrodes (Fig 6I), which showed that the aperiodic measures were mostly strongly correlated with each other.

We next examined a second, larger EEG dataset (Fig 7A-B), both as a replication of the first dataset, and to investigate the relationship between measures of aperiodic activity and age, as an example feature that measures of aperiodic activity are often compared to. We again computed measures of the aperiodic exponent, using both *specparam* and *irasa* (Fig 7C-D), finding a high correlation, computed both across subjects (r=0.950 [0.919-0.966], p<0.001) and across electrodes (r=0.973 [0.955-0.981], p<0.001), as well as computing the dominant oscillatory activity (Fig 7E). We also again computed the correlations between the aperiodic measures, as well as to the periodic measure, across subjects (Fig 7H) and across electrodes (Fig 7I), which again supported typically strong correlations across aperiodic measures, mimicking the results in the first EEG dataset.

**Figure 7).**
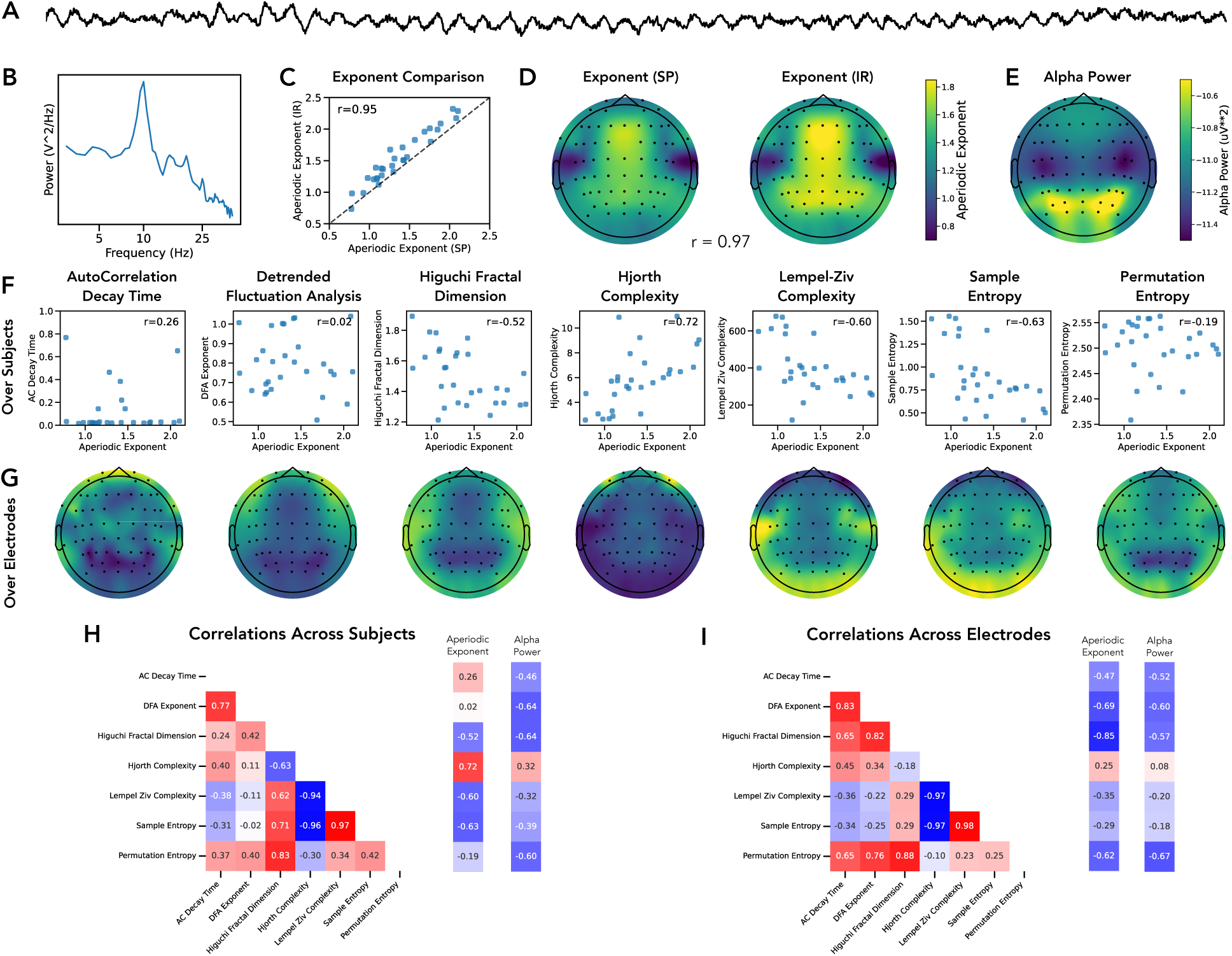
Application of aperiodic measures to an EEG dataset. **A)** An example time series from the EEG dataset. B) An example power spectrum from the EEG dataset (same data as **A**). **C)** Across subject comparison of aperiodic exponent estimations, comparing the specparam (SP) and irasa (IR) methods. **D)** Across electrode comparison of aperiodic exponent estimations. **E)** Topography of alpha power in the dataset. **F)** Across subject comparisons of time domain aperiodic measures, comparing each to the measured aperiodic exponent. **G)** Topographies of the aperiodic measures. **H)** Across subject correlations of the aperiodic measures. **I)** Across electrode correlations of the aperiodic measures.

We also computed the relationship between the spectral exponent, as measured by *specparam*, and age of the participants, replicating a strong association between the two (r=-0.519 [-0637--0.374], p<0.001). We also computed the time domain measures of aperiodic activity and computed each of their correlations with age (Fig 7F). We then sought to evaluate to what extent each measure is independently correlated with age, as compared to what extent the relationship to age reflects shared variance across multiple aperiodic measures. To do so, we recomputed the semi-partial correlations between each measure and age, after removing the impact of the aperiodic exponent (Fig 7G). We then computed the differences between the original correlations, and the semi-partial correlations, which showed that each measure had a significantly different correlation value after removing the aperiodic exponent (all p’s < 0.001), all of which reflected a decrease in magnitude of the correlation, except for the autocorrelation decay time. This suggests that overall, the relationship between aperiodic measures and age reflects shared variance across the difference measures, except for the autocorrelation decay time.

Finally, we examined an openly available dataset of intracranial EEG data (Frauscher et al., 2018), which permitted the examination of potential modality-specific differences (Fig 8A-B). In this dataset, based on the observation of ‘knees’ in the power spectra, and consistent with recent work in intracranial recordings showing that aperiodic activity in intracranial activity tends to have such a knee (Gao et al., 2020), we fit spectral measures using a model with a knee across the broader frequency range of 1-60 Hz. We fit both *specparam* (Fig 8C) and *irasa* (Fig 8D) methods and compared the results for both the aperiodic exponent (Fig 8E, r=0.862 [0.836-0.885], p=<0.001, median difference: 0.30), and the aperiodic knee (Fig 8F, r=0.908 [0.890-0.923], p=<0.001, median difference: 1.62). While there are no ground truth measures available for an empirical dataset, based on the presence of prominent knees and large bandwidth peaks (Figure 8B-C), the results of the simulations suggests that *specparam* is likely to have outperformed *irasa* in such situations. We also computed the set of aperiodic measures and computed the correlations across all measures, as well as to the aperiodic exponent, aperiodic knee, and dominant peak power (Fig 8H). Notable relationships here include that the aperiodic knee is highly correlated with the autocorrelation decay time, as expected from their theoretical relationship (Gao et al., 2020), and also very correlated with Hjorth complexity.

**Figure 8).**
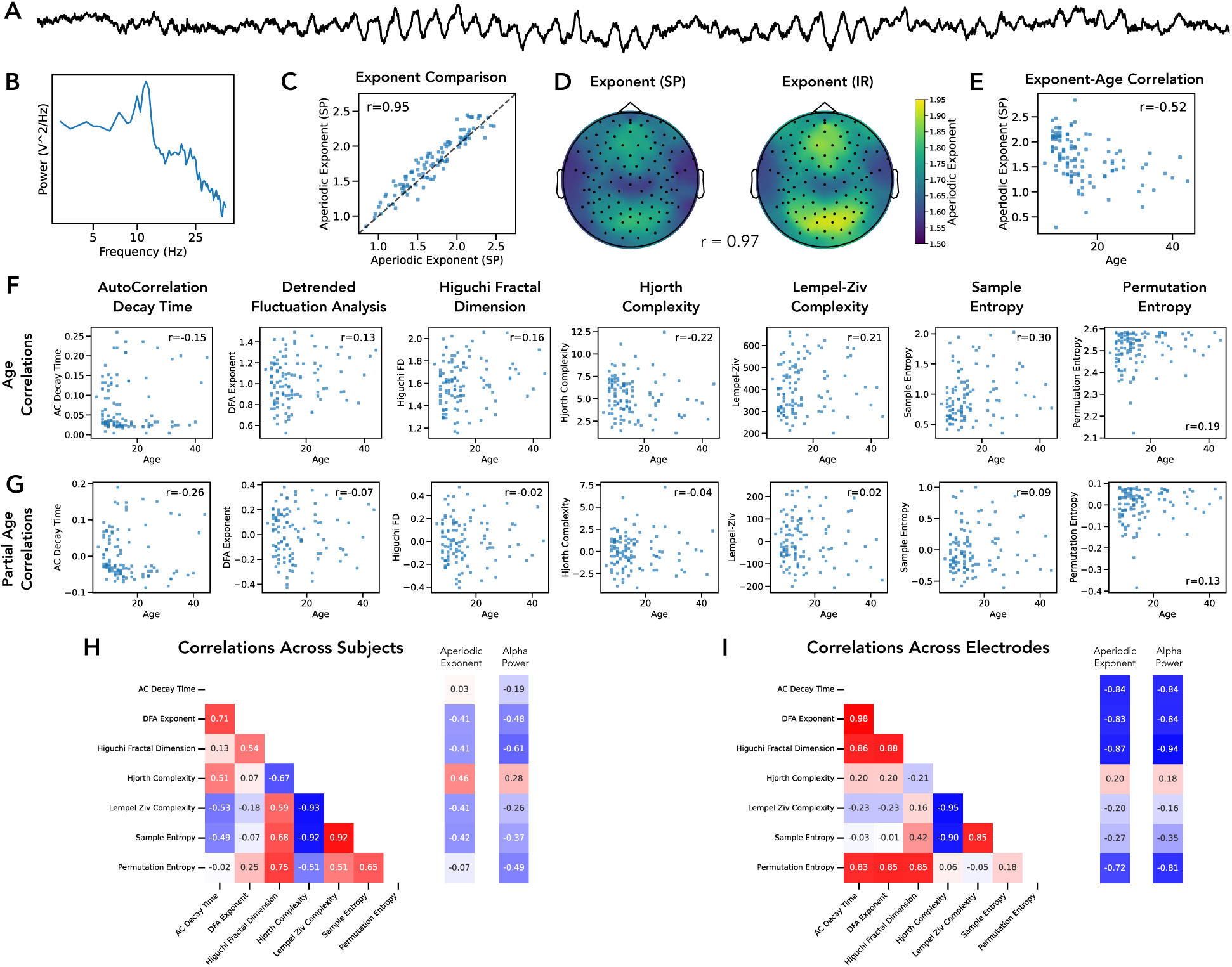
Application of aperiodic measures to a developmental EEG dataset. **A)** An example time series from the developmental EEG dataset. **B)** An example power spectrum from the EEG dataset (same data as **A**). **C)** Across subject comparison of aperiodic exponent estimations, comparing the *specparam* (SP) and *irasa* (IR) methods. **D)** Across electrode comparison of aperiodic exponent estimations. **E)** Across subject comparison of the relationship between estimated aperiodic exponent (at electrode Cz) and age across the whole dataset. **F)** Age correlations showing the across subject comparison of the relationship between time domain aperiodic measures and age. **G)** Partial age correlations showing the across subject comparison of the relationship between time domain aperiodic measures, after removing the covariance with the aperiodic exponent and age. **H)** Across subject correlations of the aperiodic measures. **I)** Across electrode correlations of the aperiodic measures.

**Figure 9).**
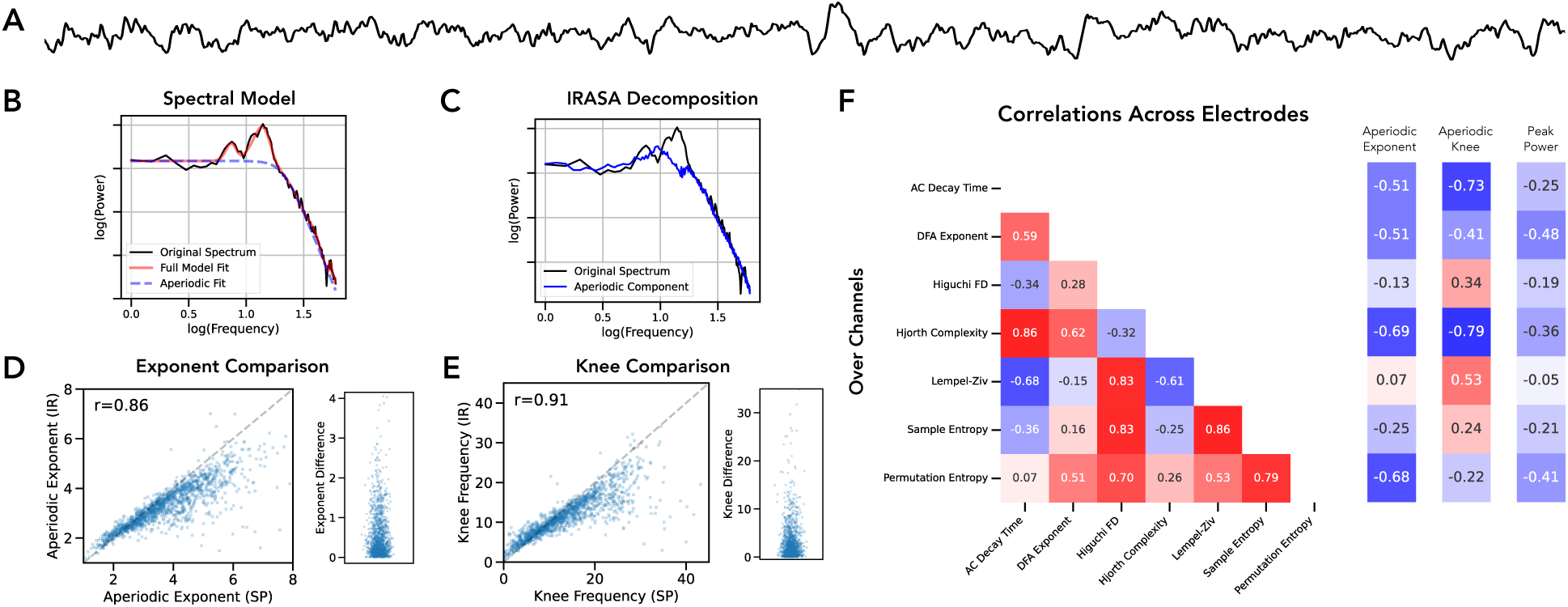
Application of aperiodic measures to an iEEG dataset. **A)** An example time series from the iEEG dataset. **B)** An example spectral model from the *specparam* method. **D)** An example spectral decomposition from the irasa method. D) Across electrode comparison of aperiodic exponent estimations, comparing the *specparam* (SP) and *irasa* (IR) methods (left) and differences between measures (right). **E)** Across electrode comparison of aperiodic knee estimations, comparing the *specparam* and *irasa* methods (left) and differences between measures (right). **F)** Across electrode correlations of the aperiodic measures.

Collectively the literature, simulation, and empirical analyses here suggest some key themes across the range of examined methods. To a first approximation, the pattern of results in the simulation analyses suggests that all the included methods are highly correlated, and that the dominant feature driving this similarity is the aperiodic activity. Notably, there are idiosyncrasies to how the different methods relate to periodic activity, including that the frequency domain methods are largely invariant to periodic activity (by design), while the time domain measures are generally much more sensitive to oscillatory features. For frequency domain methods, *specparam* outperforms simpler fitting methods, and comparing *specparam* and *irasa* suggests they are comparable in situations with a single 1/f regime and narrowband peaks, with *specparam* generalizing better to situations with knees in the aperiodic component and/or high-bandwidth oscillatory components. The patterns of results established in the simulated data are largely replicated in the empirical datasets. The empirical datasets also allowed for an example empirical analysis relating the aperiodic methods to age and showing this largely reflets shared variance. Overall, while the idiosyncrasies across comparisons preclude simplistic 1-to-1 mappings between measures, the overall pattern of results is consistent with the original hypothesis that across distributed and often disconnected literatures using these different methods, the results likely reflect overlapping patterns in the data, such that much can potentially be inferred about aperiodic neural activity, it’s correlates, and potential interpretations by examining across this literature.

## 4. Discussion

In this project we sought to collect, examine, and compare the broad set of methods that have been employed in the study of ‘aperiodic’ activity, broadly construed, in neuro-electrophysiological recordings. In evaluating the previous literature, we find a rich history, albeit one split across different literatures with idiosyncratic methods and interpretations. There is also a notable recent increase in interest in aperiodic neural activity, consistent with contemporary discussions on methodological and conceptual motivations for studying aperiodic activity (Donoghue & Watrous, 2023; Waschke, Kloosterman, et al., 2021). This approach highlights several key factors that are salient for considering the study of aperiodic neural activity, including that 1) there are a numerous methods employed across a large number of studies suggesting a considerable amount of information in literature on this topic, albeit fractured across multiple distinct areas, 2) there is a high degree of similarity in these methods, suggesting that overall they capture similar and overlapping features of the data, with the caveat that 3) there is also a notable degree of idiosyncrasy across the different approaches (including non-linear relationships between methods and differences in what features of the data they are sensitive to), such that for any given pairwise comparison or relationship to covariates of interest there are nuances in terms of how they relate to each other.

In evaluating a large collection of time domain methods, we find varied patterns of relationships within and between methods. A notable property of all the time-domain measures examined here (as compared to the frequency domain methods) is that they do not separate periodic and aperiodic components in the data. Due to this, while measures such as *dfa* and fractal dimension do approximate the expected result on purely aperiodic signals, in the context of combined signals with oscillatory components, the measured results may not specifically reflect the aperiodic activity in the data, and as such violate the theoretical expectations of such measures. Relatedly, when comparing these methods to frequency domain methods, these methods are not, by design, selective to the same properties of the data. Based on the simulated data, in which correlations between methods were systematically higher in aperiodic (as compared to combined) signals, one key takeaway is that the extent to which these methods differ may not relate primarily to differences in their sensitivity to aperiodic activity, but rather due to differences in their sensitivity to periodic activity.

Notably, most of the time domain methods employed here were developed in different fields, and then adopted into neuroscience, and as such it is important to consider to what extent the properties of neural data are consistent with the context(s) in which the method(s) were developed, and their underlying assumptions of the data. Overall, this highlights that if the goal of a particular analysis is to characterize the complexity (broadly construed) of a signal, abstracted across the different features of the data that may contribute to a measure, then the time domain measures may be useful. However, in so far as the goal is to specifically quantify aperiodic components (distinct from periodic activity), then such measures may not be appropriately specific to the desired features, and frequency domain methods may be more appropriate. An interesting avenue for future work would be to continue to examine and develop approaches for separating aperiodic and periodic components in the time domain (Samaha & Cohen, 2022), such that time domain methods could be applied to isolated components.

For frequency domain methods that measure aperiodic activity, we first compared different approaches for directly fitting the aperiodic exponent from power spectra and found that spectral parameterization (*specparam*) has the lowest average error, and lowest variance, outperforming the other spectral fitting approaches. This supports that explicitly and jointly modeling both periodic and aperiodic components is a beneficial approach (Donoghue, Haller, et al., 2020). Comparing *specparam* to the *irasa* method, we find that both methods are performant with similar results in applications in which the main goal is to fit a single aperiodic exponent, for example in EEG data. However, in the presences of large bandwidth peaks and/or aperiodic knees, the performance between the two methods starts to diverge, and based on simulations, the *irasa* method has lower accuracy in such cases – consistent with assumptions and known limitations of *irasa* (Gerster et al., 2022; Wen & Liu, 2016). This pattern of findings is consistent with other work that has compared between these methods, finding a generally high consistency for exponent estimations (Ouyang et al., 2020), with notable differences in the presence of a knee (Donoghue, Haller, et al., 2020). Overall, of the examined frequency domain methods, *specparam* has the advantage of supporting multiple different fit approaches that can best capture the data across difference contexts, being robust to both periodic and aperiodic variations.

Also highlighted by the spectral parameterization method is the importance of decisions such as the frequency range and model form to fit to. While these decisions are made explicit in methods that require the definition of such settings in the frequency domain, we note that such considerations are also salient in time domain methods, though often implicitly. Preprocessing steps that include filtering the data may significantly impact measured results, analogous to choosing different frequency ranges in the frequency domain. In addition, choosing a particular time domain method can be considered as the selection of a model – and different variants (or model forms) may be available. For example, while we here focused on ‘single timescale’ time domain methods / models, there are multiscale (multifractal) variants of time domain methods, including of *dfa* (Kantelhardt et al., 2002) and entropy (Costa et al., 2002). Such multiscale time domain methods are somewhat analogous to the use of ‘knee’ models in the frequency domain. While this study did not focus on such multifractal situations, in the iEEG data analyses (in which knees are common), we did fit spectral knee models, with the resultant spectral features still being highly correlated with the single-scale time domain methods. Future work should include further investigation of multi-timescale (multi-fractal) methods, in particular in the time domain, investigating how these methods relate to neural data. In doing so, we can continue to develop best-practice guidelines for developing measurement approaches that explicitly test method and model assumptions and evaluate their performance on empirical data.

In this work, we extend the analyses of previous investigations that have compared different methods for measuring aperiodic activity. Such investigations have typically investigated two or three methods at a time without the use of ground-truth simulated data that mimic neural recordings. For example, a strong correlation between measures of the fractal dimension (as measured by the correlation dimension) and the aperiodic exponent has been reported (Krakovská & Štolc, 2008). Similarly, previous work has reported strong correlations between the aperiodic exponent and Lempel-Ziv complexity (Alnes et al., 2023; Medel et al., 2023), as well as between the aperiodic exponent and various measures of entropy (Kosciessa, Kloosterman, et al., 2020; Miskovic et al., 2019; Waschke et al., 2017). Recently there have also been other broader investigations, including an investigation of over 100 complexity measures (Makowski et al., 2022), finding a high degree of shared variance across them, and similarly a comparison of thousands of different time-series measures in MEG data found that the first principal component across all methods largely reflected the structure of the power spectrum (Shafiei et al., 2023). Collectively, the high degree of correlation between measures in this report is consistent with other related investigations, whereby we contribute here a specifically curated analysis of measures for aperiodic activity from across broad literatures, against ground truth simulated data that matches the properties of neuro-electrophysiological time series.

Given the high degree of similarity across the different measures, a key question is whether different studies employing different methods (and associated theories) and finding seemingly related results (for example, that complexity decreases and aperiodic exponent increases during sleep) reflect different quantifications of the same underlying changes in the data, and/or reflect independent variance. The main set of results here, showing high correlations across the methods in simulated and empirical data, is consistent with these methods capturing shared variance, though does not definitively establish the degree to which such measures covary with covariates of interest. In an example analysis in which we examined the relationship between aperiodic measures and age, we found that by regressing out the shared variance of the aperiodic exponent, all the examined time domain methods (except autocorrelation) were significantly reduced in their correlation with age. This is consistent with them all capturing, at least in part, the same variance in the data. Overall, this project suggests that many previous papers are potentially measuring and reporting the same underlying effects with different measures (and potentially different interpretations). However, we also note that there are significant idiosyncrasies and non-linearities in what the measures are sensitive to and how they covary, such that for any set of measures and/or covariates, a definitive answer requires follow up work to evaluate the specific measures and relationships under study.

Another key consideration is what these empirical relationships between the methods means in relation to the different terminology, interpretations, and conceptualizations related to the different approaches. Notably, while this study has used the term ‘aperiodic’, chosen to be a theoretically neutral and descriptively broad term to refer to a range of related ideas and analyses in the literature, this term itself is by no means a universal term in the relevant literature, and may not fully capture the range of ideas and interpretations across the literature. Some work uses the term ‘noise’ (or ‘neural noise’), which may be descriptively relevant in terms of such activity being appropriately described ‘colored noise’, however this can also be counterproductive if examining such activity as salient, physiological, and informative component of the signal. The term ‘complexity’ is also common, not only in relation to ‘complexity’ measures but also commonly in reports on entropy and fractal measures, and is a useful general term – however, it is worth noting that there is no singular agreed upon definition of complexity (related to why different measures of complexity can give opposing answers). Overall, we find there is no singular term that is necessarily preferred or most appropriate, but emphasize the utility of connecting and comparing across literature that uses different terminology.

Related to the variation in terminology, aperiodic neural activity has been analyzed under multiple conceptual frameworks and associated interpretations. Across the literature, findings have also been interpreted and contextualized in various overlapping ways – for example, measures of entropy / complexity have been interpreted in relation to neural variability (Waschke, Kloosterman, et al., 2021); fluctuation and fractal measures have been interpreted by focusing on fractal properties and self-similarity (Eke et al., 2002; Schaefer et al., 2014) and/or in relation to considering the underlying systems as nonlinear dynamical systems and concepts such as criticality (Le Van Quyen et al., 2003; Stam, 2005); and measures of the spectral exponent have been interpreted as functional interpretations, such as relating to ‘neural noise’ (Voytek et al., 2015), physiological models and interpretations, such as relating aperiodic activity to post-synaptic potentials and/or excitatory-inhibitory balance (Freeman & Zhai, 2009; Gao et al., 2017; Miller et al., 2009) and theoretical interpretations, such as with appeal to the scale-free properties (Grosu et al., 2023; He, 2014). Notably, the conceptualizations, interpretations, and implications across these theoretical positions are not strictly distinct, with much potential overlap – though fully understanding which findings and interpretations are (in)consistent with each requires both a clear mapping between the empirical findings, and a careful comparison of their potential interpretations.

A full comparison of the conceptual frameworks and interpretations is out of scope of this paper. However, this project emphasizes the degree to which a single method and associated terminology that appears to fit with the data could easily be remeasured and redescribed with an alternative approach that could seem to imply different interpretations. To what extent such conceptualizations are consistent and coherent, and/or potentially mutually exclusive should be topic of future work systematically comparing the theoretical models, especially given the large overlap in methods and results demonstrated here. In relation to this, we re-emphasize a point made by several papers discussing methods (Eke et al., 2002; Le Van Quyen et al., 2003), which is that these technically precise terms and theories – such as referring to ‘scalefree’ or ‘fractal’ properties of the data, or to ‘critical’ or ‘dynamical’ systems – have assumptions and precise definitions that need to explicitly tested in the data, and if/when such assumptions are violated the methods and/or interpretations cannot be applied to such situations. For a simple example, discussions of ‘scalefree’ activity may be inaccurate in situations in which the self-similarity, 1/f property does not extend over a sufficient range (for example, if there is a knee). By combining explicit definitions and testing of the different theories assumptions and predictions, in combination with an understanding of the differences and similarities across these related measures, future work can seek to map out the similarities and differences across these conceptual frameworks in order to come to a better understanding of nature and interpretations of aperiodic neural activity.

The key goal of this project was to evaluate the relationships between different methods, and not to make any recommendations on the ‘best’ methods – nevertheless, some general recommendations can be made. The first is that, for research topics of interest, it may be useful to explore the literature for studies that use different analysis methods and even different language but may ultimately reflect the same patterns of interest in the data under study. In designing analyses, it’s important to consider what measure assumptions / requirements, and what features a method is sensitive to. For example, the spectral domain methods employed here explicitly seek to separate aperiodic and periodic activity, whereas, broadly speaking, the time domain methods reflect a combined measure across both, with varying sensitivity to different features. Depending on the goals of the study, different methods may be more or less suited for the analysis. For example, time domain complexity measures may reflect oscillatory activity in ways that is undesired in certain contexts, and/or for frequency domain measures features such as large bandwidth peaks and/or knees may suggest the use of *specparam* over *irasa*. In addition, we emphasize that simulation-based evaluations and/or direct comparison of different measures can be productive in evaluating what features drive a measured change and how different methods compare to each other.

## 5. Conclusion

Aperiodic electrophysiological neural activity is a prominent and dynamic component of neural field recordings with many known cognitive, behavioral, and disease correlates. Despite this, there is currently no consensus for best practices or comparisons across approaches for quantifying this signal. This is complicated by the many conceptual frameworks that determine the analytical approach used. Here, we start with a literature analysis to identify commonly applied methods, and subsequently used a simulation-driven approach to systematically evaluate and compare this set of methods. Overall, we find a high degree of similarity between the results of these methods, in simulated and empirical datasets, suggesting that they likely capture the same or highly overlapping properties of the data. We suggest that future work should continue to investigate the nuances of the relationships between these methods, and systematically explore how this relates to theories and conceptualizations of aperiodic neural activity.

## Acknowledgements

We would like to thank members of the Voytek lab, past and present, for insightful discussions, as well as all the contributors to the open-source software that was used by this project.

## Abbreviations

EEG: electroencephalography
iEEG: intracranial EEG
MEG: magnetoencephalography
LFP: local field potential
DSP: digital signal processing
*dfa*: detrended fluctuation analysis
*specparam*: spectral parameterization
*irasa*: irregular resampling auto-spectral analysis

## Materials Descriptions & Availability Statements

### Project Repository

This project is also made openly available through an online project repository in which the code and data are made available, with step-by-step guides through the analyses.

Project Repository: http://github.com/AperiodicMethods/AperiodicMethods

### Project Website

This project is also hosted on an openly available project website. The project website is created by converting the notebooks used in this project to create static HTML pages, showing the code and outputs, including all the figures from this project, as well as notes and comments.

Project Website: https://aperiodicmethods.github.io/

### Data

This project uses literature data, simulated data, and empirical EEG and iEEG datasets. The literature data was collected from the Pubmed database, with the code to reproduce available in the project repository. The simulations used in this project are created with openly available software packages. Settings and code to re-generate simulated data, as well as copies of the simulated data that were used in this investigation, are available in the project repository. The EEG data includes a dataset collected in the VoytekLab at UC San Diego, as well as the open-access ChildMind MIPDB dataset. The iEEG data is open-access data from the MNI database.

### Code

Code used and written for this project was written in the Python programming language. All the code used within this project is deposited in the project repository and is made openly available and licensed for re-use.

All code used in this project is available at:

https://github.com/AperiodicMethods/AperiodicMethods

Open-source tools used in this project:

lisc https://github.com/lisc-tools/lisc
neurodsp https://github.com/neurodsp-tools/neurodsp
specparam https://github.com/fooof-tools/fooof
antropy https://github.com/raphaelvallat/antropy
neurokit2 https://github.com/neuropsychology/NeuroKit

